# Hippocampal place cell sequences are impaired in a rat model of Fragile X Syndrome

**DOI:** 10.1101/2024.10.18.619112

**Authors:** Margaret M. Donahue, Emma Robson, Laura Lee Colgin

**Affiliations:** Center for Learning and Memory, The University of Texas at Austin, Austin, TX 78712; Institute for Neuroscience, The University of Texas at Austin, Austin, TX 78712; Department of Neuroscience, The University of Texas at Austin, Austin, TX 78712

## Abstract

Fragile X Syndrome (FXS) is a neurodevelopmental disorder that can cause impairments in spatial cognition and memory. The hippocampus is thought to support spatial cognition through the activity of place cells, neurons with spatial receptive fields. Coordinated firing of place cell populations is organized by different oscillatory patterns in the hippocampus during specific behavioral states. Theta rhythms organize place cell populations during awake exploration. Sharp wave-ripples organize place cell population reactivation during waking rest. Here, we examined the coordination of CA1 place cell populations during active behavior and subsequent rest in a rat model of FXS (*Fmr1* knockout rats). While the organization of individual place cells by the theta rhythm was normal, the coordinated activation of sequences of place cells during individual theta cycles was impaired in *Fmr1* knockout rats. Further, the subsequent replay of place cell sequences was impaired during waking rest following active exploration. Together, these results expand our understanding of how genetic modifications that model those observed in FXS affect hippocampal physiology and suggest a potential mechanism underlying impaired spatial cognition in FXS.

**Significance Statement:** Fragile X Syndrome (FXS) is a neurodevelopmental disorder that can cause impaired memory and atypical spatial behaviors such as “elopement” (i.e., wandering off and becoming lost). Activity in the CA1 subregion of the hippocampus supports spatial memory and spatial cognition, making it an important candidate to study in the context of FXS; however, how neuronal population activity in CA1 is affected by FXS is poorly understood. In this study, we found that the coordination of populations of CA1 neurons during active behavior and waking rest was impaired in a rat model of FXS. These results reveal hippocampal physiological deficits that may contribute to cognitive impairments in FXS.

## Introduction

Fragile X Syndrome (FXS) is a widespread neurodevelopmental disorder that is caused by epigenetic suppression of the X-chromosome linked *Fmr1* gene and subsequent loss of Fragile X Messenger Ribonucleoprotein (FMRP) (De Boulle et al., 1993; Kremer et al., 1991; Mikiko et al., 1995; Pieretti et al., 1991). Patients with FXS are impaired in memory tasks (Cornish et al., 1998; Gallagher & Hallahan, 2012; Jäkälä et al., 1997) and often display elopement behaviors (Machalicek et al., 2014; Muller et al., 2019). Mouse and rat models of FXS (“FXS mice” or “FXS rats”, respectively) have been produced by knocking out the *Fmr1* gene (Bakker et al., 1994; Tian et al., 2017). Spatial memory deficits have been reported for both FXS mice and FXS rats (Asiminas et al., 2019; Bakker et al., 1994; Mineur et al., 2002; Tian et al., 2017; Till et al., 2015; Van Dam et al., 2000). FMRP is an RNA-binding protein (Siomi et al., 1993) that interacts with many neuronal proteins (Darnell et al., 2011; for a review, see Darnell & Klann, 2013) and is highly expressed in the hippocampus (Ludwig et al., 2014).

The hippocampus is thought to support memory and spatial cognition through the activity of place cells, neurons with spatial receptive fields known as “place fields” (O’Keefe, 1976; O’Keefe & Dostrovsky, 1971). Populations of place cells form sequences that represent trajectories through an environment. These sequential firing patterns are coordinated by hippocampal rhythms that are differentially associated with behavioral states. During active exploration of an environment, place cells are coordinated by the theta rhythm, a ∼6-10 Hz oscillation occurring prominently in local field potential (LFP) recordings from the hippocampus (Buzsáki, 2002). As an animal moves through a cell’s place field, place cells fire at progressively earlier phases of the theta rhythm in a phenomenon known as “theta phase precession” (O’Keefe & Recce, 1993). Coordinated theta phase precession across multiple place cells with adjacent place fields would be expected to result in an organized sequence of spikes within an individual theta cycle that represent a rat’s previous, current, and future locations. Such sequential activation of place cells within a single theta cycle has been observed in rats and termed a “theta sequence” (Foster & Wilson, 2007). Previous work has suggested that hippocampal networks are hypersynchronized in FXS (Arbab, Battaglia, et al., 2018; Talbot et al., 2018), which could impair organization of cells in theta sequences. However, the activity of theta-coordinated sequences of place cells is yet to be investigated in a rodent model of FXS.

Sequences of place cells that were activated during active exploration of an environment are reactivated or “replayed” in a time-compressed manner during subsequent waking rest and non-REM sleep (Kudrimoti et al., 1999; Lee & Wilson, 2002). Place cell replay co-occurs with characteristic events in the hippocampal LFP called sharp wave-ripples (Kudrimoti et al., 1999; Lee & Wilson, 2002; Nádasdy et al., 1999). Sharp wave-ripples during sleep are abnormal in FXS mice (Boone et al., 2018), suggesting that coordination of place cell populations during sleep and rest may be disrupted in FXS. However, replay of place cell sequences has not been examined in FXS models.

Here, we examined the activity of coordinated sequences of CA1 place cells in FXS rats traversing a familiar circular track and subsequently resting. During active behaviors, we found that the coordination of individual place cells by the theta oscillation was normal in FXS rats.

However, theta-coordinated sequences of place cells in FXS rats were less temporally compressed and represented shorter paths than those in wild-type (WT) control rats. Further, replay of place cell sequences during rest was abnormally slow in FXS rats, with replay events exhibiting longer durations and representing less temporally compressed spatial trajectories than in WT rats. These findings raise the possibility that the abnormal coordination of sequences of hippocampal place cells may contribute to impairments in spatial memory and cognition in FXS.

## Materials and Methods

Additional data collected from one of the rats (rat 418) used in this study was presented in a previous study (Robson et al., 2024). Surgery, data acquisition, histology, and spike sorting methods in this paper are identical to methods described in that study and re-stated below.

### Subjects

Twelve male Sprague-Dawley rats (Inotiv) were used for this study. Six rats were *Fmr1*-knockout rats (*SD-Fmr1-nulltm1Sage*), and six were littermate WT control rats. As FXS is an X-chromosome linked disorder, FXS has a higher prevalence and increased symptom severity in males. Thus, we chose to use male rats for this study. Rats were between the ages of 3-11 months old at the time of surgery. Prior to surgery, rats were double-or triple-housed in genotype-matched groups. After surgery, rats were singly housed in custom-built acrylic cages (40 cm x 40 cm x 40 cm) containing enrichment material (wooden blocks, paper towel rolls, etc.) and maintained on a reverse light cycle (light: 8 p.m. – 8 a.m.). Rats were housed next to their former cage mates after recovering from surgery and throughout behavioral testing. Rats recovered from surgery for at least one week before behavioral training resumed. All behavioral experiments were performed during the dark cycle. When necessary to encourage spatial exploration, one rat (rat 316) was placed on a food-deprivation regimen. While on the regimen, this rat maintained ∼98% of his free-feeding body weight. Following the completion of all recording experiments, a small piece of ear tissue was collected from each rat to verify genotype. All experiments were conducted according to the guidelines of the United States National Institutes of Health Guide for the Care and Use of Laboratory Animals and under a protocol approved by the University of Texas at Austin Institutional Animal Care and Use Committee.

### Surgery and tetrode positioning

“Hyperdrives” with 14 independently movable tetrodes were implanted in eight of the rats. Hyperdrives with 21 independently movable tetrodes were implanted in four of the rats. Implants were positioned above the right dorsal hippocampus (anterior-posterior –3.8 mm from bregma, medial-lateral –3.0 mm from bregma). To implant and stabilize the hyperdrives, eleven bone screws were affixed to the skull, and the base of the implant and the screws were covered in dental acrylic. Two of the screws were connected to the recording drive ground. Prior to surgery, tetrodes were built from 17 μm polyimide-coated platinum-iridium (90/10%) wire (California Fine Wire, Grover Beach, California). The tips of tetrodes designated for single-unit recording were plated with platinum to reduce single-channel impedances to ∼150 to 300 kOhms. All tetrodes were lowered ∼1 mm on the day of surgery. Thereafter, tetrodes were slowly lowered to the hippocampal pyramidal cell body layer over the course of several weeks except for one tetrode that was designated for use as a reference for differential recording. This reference tetrode was placed in an electrically quiet area approximately 1 mm above the hippocampus and adjusted as needed to ensure quiescence. All four wires of this tetrode were connected to a single channel on the electrode interface board. The reference signal was duplicated using an MDR-50 breakout board (Neuralynx, Bozeman, Montana) and recorded continuously to ensure that unit activity or volume conducted signals of interest were not detected. Another tetrode was placed in the apical dendritic layer of CA1 to monitor LFPs and guide placement of the other tetrodes using estimated depth and electrophysiological hallmarks of the hippocampus (e.g., sharp wave-ripples).

### Data acquisition

Data were acquired using a Digital Lynx system and Cheetah recording software (Neuralynx, Bozeman, Montana). The recording setup has been described in detail previously (Hsiao et al., 2016; Zheng et al., 2016). Briefly, LFP signals from one randomly chosen channel per tetrode were continuously recorded at a 2000 Hz sampling rate and filtered in the 0.1–500 Hz band. Input amplitude ranges were adjusted before each recording session to maximize resolution without signal saturation. Input ranges for LFPs generally fell within ±2,000 to ±3,000 μV. To detect unit activity, all four channels within each tetrode were bandpass filtered from 600 to 6,000 Hz. Spikes were detected when the filtered continuous signal on one or more of the channels exceeded a threshold of 55 µV. Detected events were acquired with a 32,000 Hz sampling rate for 1 ms. For both LFPs and unit activity, signals were recorded differentially against a dedicated reference channel (see “*Surgery and tetrode positioning*” section above).

Videos of rats’ behavior were recorded through the Neuralynx system with a resolution of 720 × 480 pixels and a frame rate of 29.97 frames/s. Rat position and head direction were tracked via an array of red and green or red and blue light-emitting diodes (LEDs) on a HS-54 or HS-27 headstage (Neuralynx, Bozeman, Montana), respectively.

### Behavior

Rats were trained to run unidirectionally on a 1-meter diameter circular track. The track was 0.5 meters in height. Rats ran four 10-minute sessions on the track per day. Inter-session rest periods were 10 minutes, and the rat rested in a towel-lined flowerpot in the recording room during rest sessions. Rats additionally rested in the pot for 10 minutes prior to the start of the first track running session and after the final session for a total of five rest sessions per day. Small pieces of sweetened cereal or cookies were placed at one or two locations on the circular track to encourage running. The reward locations were kept consistent within each day but changed daily in order to prevent accumulation of place fields at the reward site (Hollup et al., 2001). To ensure that rats were familiarized with the environment prior to recording, rats completed a minimum of 2 days of the full recording session (i.e., all five rest sessions and all four run sessions) before data collection began. To compare behavior between genotypes, we calculated run speed on the circular track and number of laps completed per session. To ensure exclusion of pauses in running behavior and periods of immobility, only times when the rat was traveling at speeds greater than 5 cm/s were included when calculating run speed.

### Histology and tetrode localization

Following recording, rats were perfused with 4% paraformaldehyde solution in phosphate-buffered saline. Brains were cut coronally in 30 μm sections using a cryostat. Brain slices were immunostained for the CA2 marker Purkinje Cell Protein 4 (PCP4), allowing differentiation of all subregions of the hippocampus (CA1, CA2, and CA3). Sections were initially washed and blocked in 10% normal goat serum in TBS. Sections were incubated overnight with rabbit anti-PCP4 (1:200, Sigma-Aldrich Cat# HPA005792) diluted in TBS containing 0.05% Tween. The next day, sections were washed and incubated overnight with a secondary fluorescent antibody (Alexa Flour™-555 anti-rabbit, Thermo Fisher Scientific). All washes and incubations were performed at room temperature. Slides were mounted using DAPI Fluoromount-G (Fisher Scientific). Tetrode recording sites were identified by comparing locations across adjacent sections.

### Spike sorting

Spike sorting was performed manually using graphical cluster-cutting software (MClust, A.D. Redish, University of Minnesota, Minneapolis, Minnesota) run in MATLAB (Mathworks). Spikes were sorted using two-dimensional representations of waveform properties (i.e., energy, peak, and peak-to-valley difference) from four channels. A single unit was accepted for further analysis if the associated cluster was well isolated from, and did not share spikes with, other clusters on the same tetrode. Units were also required to have a minimum 1 ms refractory period. Units with mean firing rates above 5 Hz were considered putative interneurons and excluded from further analyses. To be included in the replay event firing analyses, a unit was required to have valid clusters in both the active exploration and rest sessions. Place cell yields for each rat for the active behavior and replay event analyses are reported in Tables 1 and 2, respectively.

**Table 1.** Total number of place cells recorded from each rat during active behavior.

| Genotype | Rat | Number of place cells |
| --- | --- | --- |
| WT | 326 ("Zachariah") | 94 |
| WT | 334 ("Chickpea") | 4 |
| WT | 335 ("Couscous") | 5 |
| WT | 392 ("Danish") | 29 |
| WT | 416 ("Desmond") | 60 |
| WT | 418 ("Hugo") | 71 |
| FXS | 316 ("Yuki") | 21 |
| FXS | 330 ("Aries") | 99 |
| FXS | 394 ("Mr. Eko") | 3 |
| FXS | 395 ("Elijah") | 36 |
| FXS | 445 ("Paddy") | 20 |
| FXS | 442 ("Pippin") | 69 |

**Table 2.** Total number of place cells recorded from each rat during replay events.

| Genotype | Rat | Number of place cells |
| --- | --- | --- |
| WT | 326 ("Zachariah") | 93 |
| WT | 392 ("Danish") | 16 |
| WT | 416 ("Desmond") | 60 |
| WT | 418 ("Hugo") | 71 |
| FXS | 316 ("Yuki") | 21 |
| FXS | 330 ("Aries") | 98 |
| FXS | 395 ("Elijah") | 18 |
| FXS | 442 ("Pippin") | 69 |

### Place cell analyses

Only cells from tetrodes in CA1 were included in analyses (see “*Histology and tetrode localization*” section above). Firing rate maps were created for each single unit using methods based on those used in our prior studies (Hwaun & Colgin, 2019; Zheng et al., 2021) (see Figure 1A-B for example rate maps). The circular track was divided into 4-degree bins. The number of spikes fired by a unit was divided by the time spent in each bin. Spikes that occurred at times when a rat was moving at speeds less than 5 cm/s were excluded. This rate map was then smoothed with a Gaussian kernel (standard deviation = 8 degrees). Rate maps were calculated individually for each run session and for all run sessions in each day. To be included for further analysis, the day-averaged rate map of a cell had to reach a minimum peak firing rate of 1 Hz.

**Figure 1.**
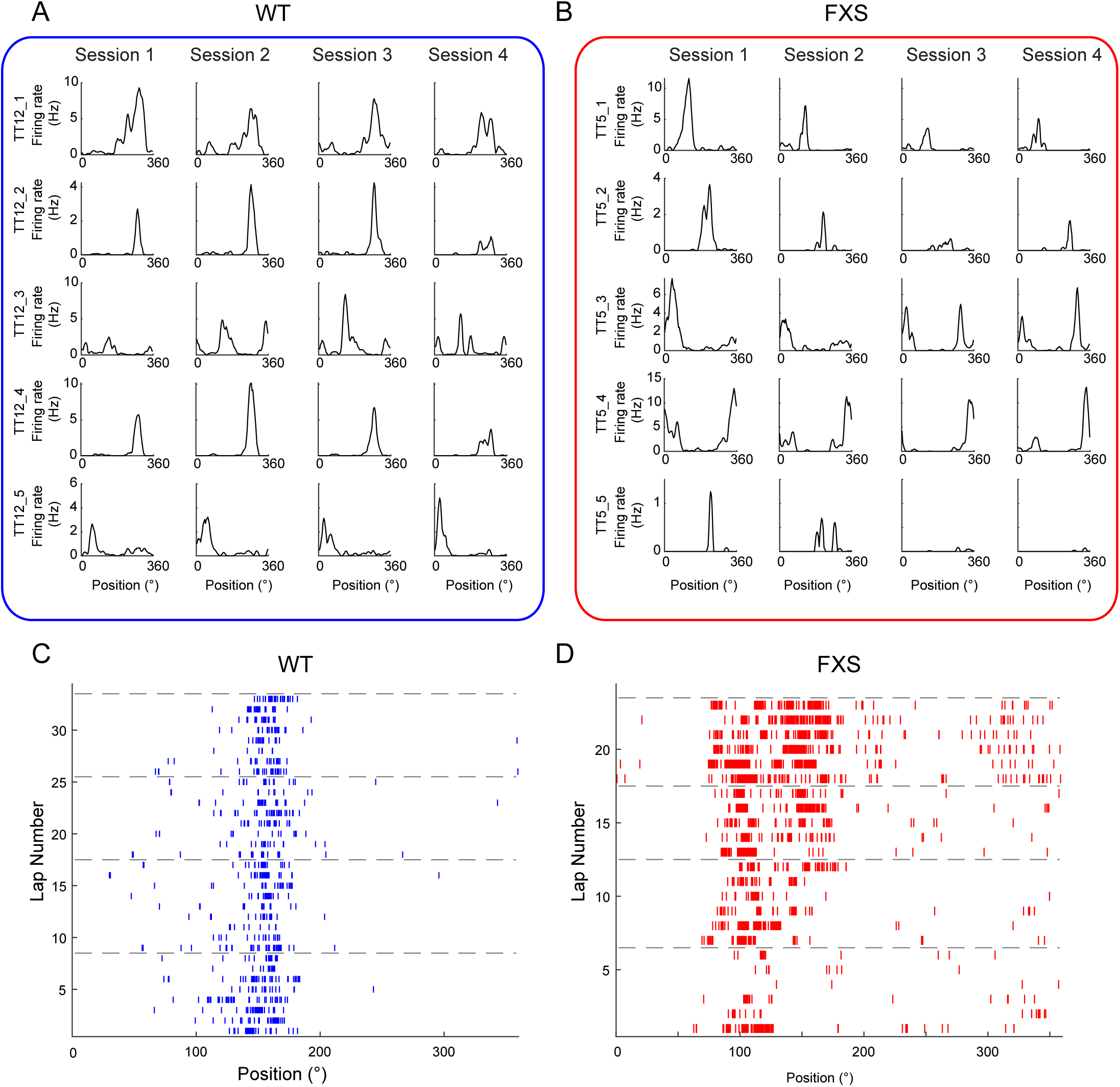
Example CA1 place cells across four recording sessions from WT and FXS rats. (A-B) Firing rates across positions on the circular track are shown for all place cells (rows) recorded from one example tetrode across all four sessions (columns) in an example day from a WT (A) and a FXS (B) rat. (C-D) Example spike raster plots for successive laps within a day for a WT (C) and a FXS (D) place cell. Dashed lines separate different sessions.

To examine stability of place cell rate maps, a Pearson correlation coefficient *R* (“spatial correlation”) was calculated for each unit between pairs of rate maps from different sessions (Figure 2A). We additionally calculated rate overlap between pairs of rate maps from different sessions for each cell to determine whether a cell’s firing rate changed significantly across sessions (Figure 2B). Rate overlap was calculated by taking the ratio of mean firing rates between two sessions, with the larger firing rate as the denominator (Colgin et al., 2010). Spatial information was calculated using the following formula (Skaggs et al., 1996):

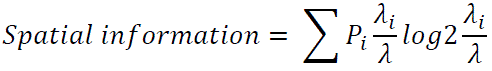

where *i* is an index of spatial bins, *P_i_* is the probability of a rat being in the *i*th bin, *λ_i_* is the mean firing rate in the *i*th bin, and *λ* is the overall mean firing rate of the cell (Figure 2C).

**Figure 2.**
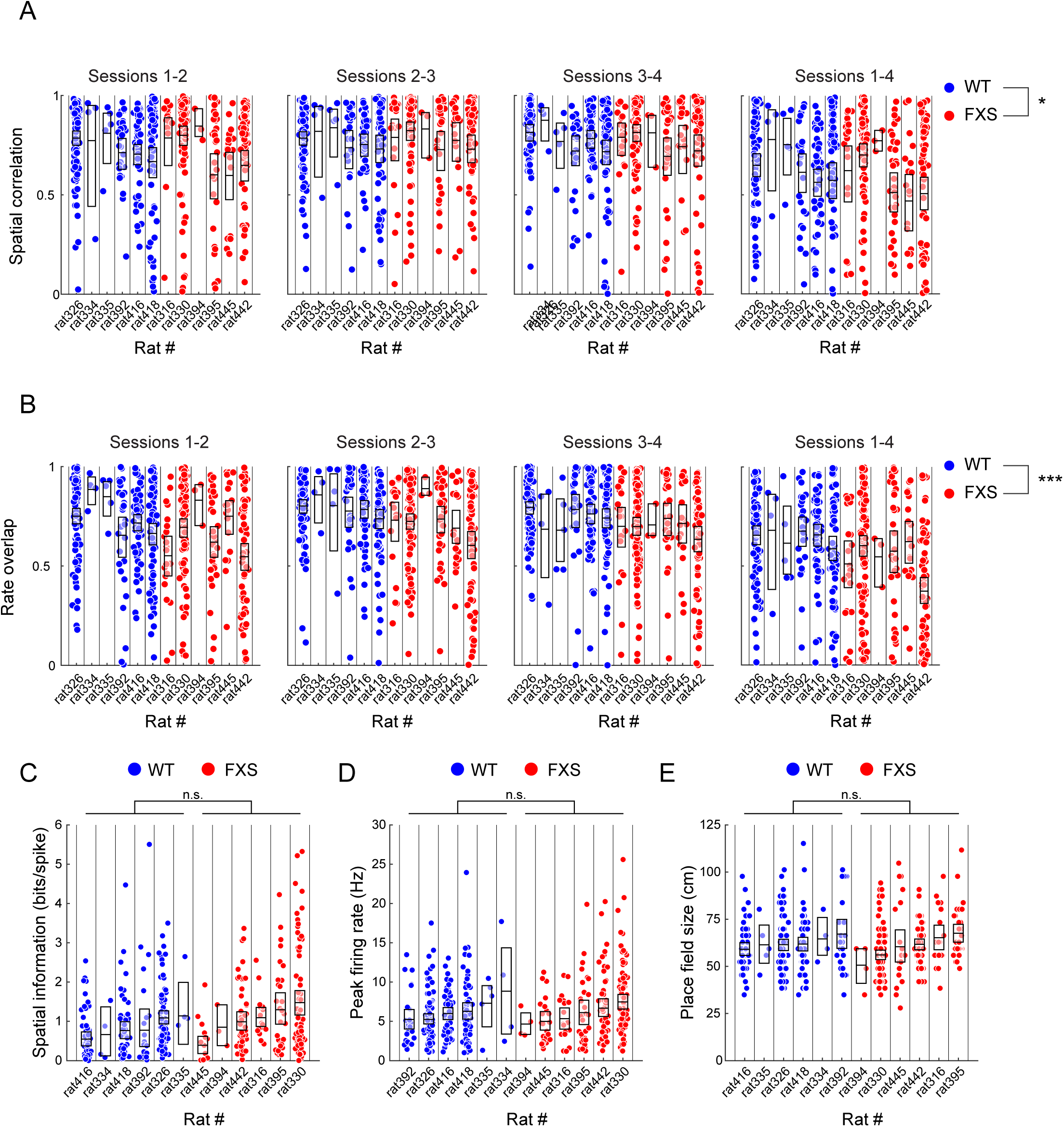
Place cell firing rate maps in FXS rats were unstable across sessions but otherwise exhibit normal place field properties. (A) Spatial correlation coefficients are shown across sessions pairs for each rat. Spatial correlation values were lower in FXS rats than WT rats, indicating impaired stability of place cell responses in FXS rats. (B) Rate overlap values are shown across session pairs for each rat. Rate overlap values were lower in FXS rats than in WT rats, indicating highly variable firing rates of place cells across sessions in FXS rats. (C) There was no difference in spatial information of place cells between WT and FXS rats. (D) Peak firing rates of place cells did not differ between WT and FXS rats. (E) Place field size did not differ between WT and FXS rats. For all plots, each dot represents a measure from one place cell. Boxes represent 95% confidence intervals of the mean for each rat.

To identify place fields (Figure 2D-E), firing rates across locations in the rate map were first z-scored. Potential place fields were identified as locations where the z-scored firing rate was greater than or equal to 2. Identified place fields were bounded by locations where z-scored firing rates fell below 0.5. To be included for further analysis, the peak firing rate in a place field had to be at least 1 Hz, and the minimum length of a place field had to be at least 18 degrees (∼15 cm).

### Phase precession analysis

Phase precession analysis was performed similarly to the analysis described in our previously published work (Bieri et al., 2014). The LFP signal from every tetrode that recorded CA1 cells identified as place cells was bandpass filtered in the theta range (i.e., between 6-10 Hz). The phase of the theta oscillation was then estimated using a Hilbert transform. Locations within each place field were normalized between 0 and 1. For each spike that a cell fired within its place field on all sessions within a day, the theta phase at the spike time was estimated using the transformed signal from the tetrode on which it was recorded. The normalized distance through the place field at the spike time was also determined. Spikes that occurred while the rat was traveling at a speed of less than 5 cm/s were excluded. A place cell had to fire a minimum of 50 spikes within its place field in order to be included in theta phase precession analysis. Circular-linear regression was then performed with theta phase as the circular variable and normalized distance through the place field as the linear variable (Figure 3A-B). The correlation coefficient for the relationship between theta phase and normalized distance (Figure 3C) was calculated using the *circ_corrcl* function from the Circular Statistics toolbox in MATLAB (Berens, 2009) (https://www.mathworks.com/matlabcentral/fileexchange/10676-circular-statistics-toolbox-directional-statistics).

**Figure 3.**
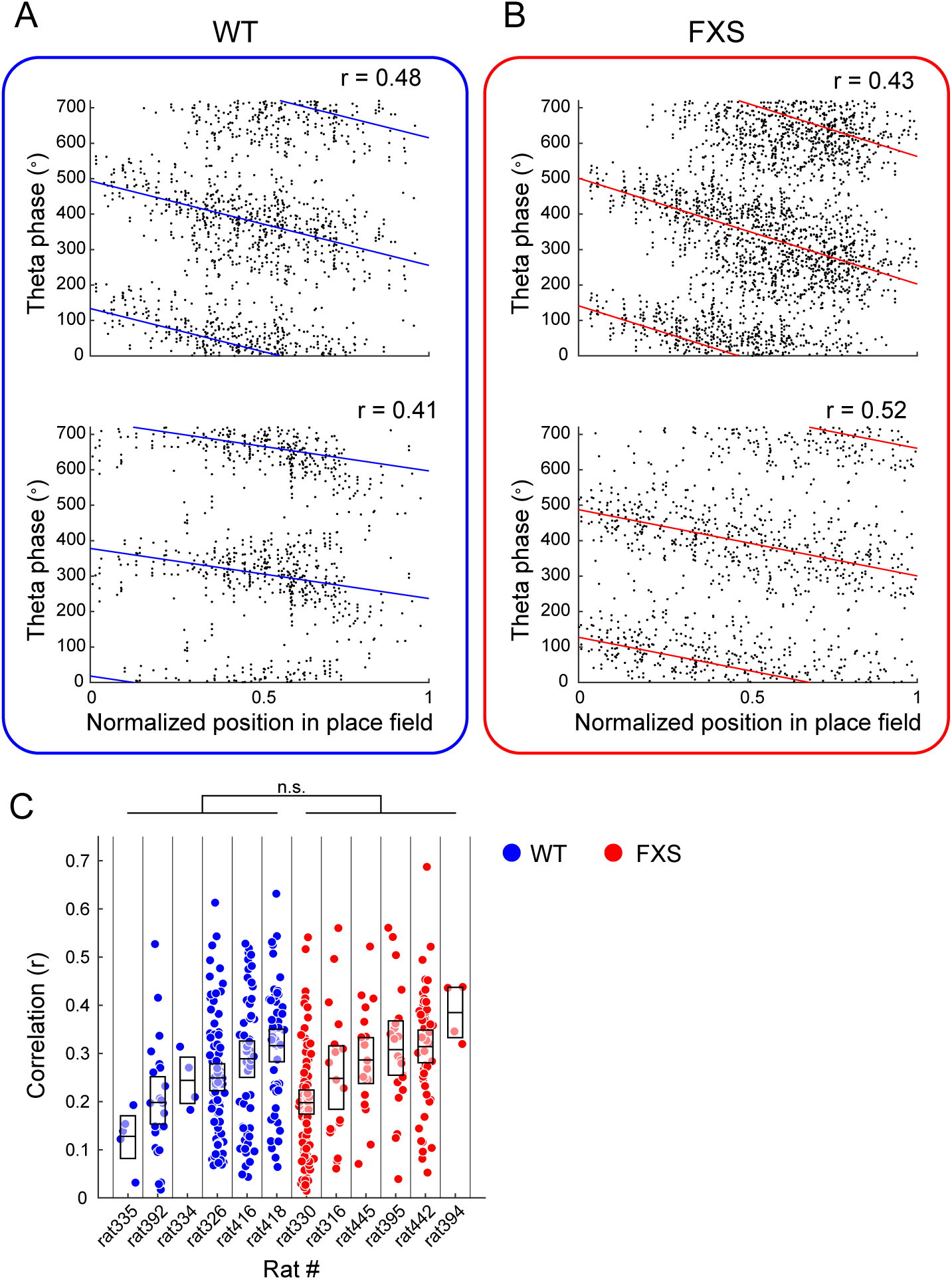
Theta phase precession was preserved in FXS rats. (A-B) Place cell phase precession plots are shown for two example place cells from (A) WT and (B) FXS rats. Each dot represents the theta phase associated with each spike and corresponding normalized position in the cell’s place field. The solid line represents the correlation between the theta phase and normalized position in place field. The magnitude of the correlation (r) is shown (top right) for each cell. (C) The correlation between theta phase and normalized position in a place field did not differ for place cells from WT and FXS rats. Each dot represents the correlation measure for one cell. Boxes represent 95% confidence intervals of the mean for each rat.

### Bayesian decoding analysis

We used a Bayesian decoding algorithm (K. Zhang et al., 1998) to estimate posterior probability distributions of angular positions represented by the spiking activity of CA1 place cell populations during track running. The probability of a rat being at position *x* given the number of spikes *n* that occurred within a given time window was defined using Bayes’ rule:

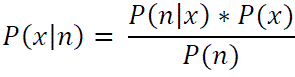

where *P(n | x)* was estimated using the averaged position tuning of each unit across all four sessions. The probability of a rat being at any given position on the circular track *P(x)* was set to 1 to avoid biasing the decoder towards any particular position on the circular track. The normalizing constant *P(n)* was set such that the posterior probability distribution *P(x | n)* summed to 1.

### Decoding accuracy analysis

To verify that a given day’s place cell yields were sufficient to accurately decode a rat’s positions on the circular track, we used a decoding accuracy analysis similar to the methods described in our previously published work (Hwaun & Colgin, 2019; Zheng et al., 2021). Positions were decoded for all times when a rat was traveling at speeds over 5 cm/s in 500 ms windows with 100 ms steps. To create confusion matrices (Figure 4A-B), the average decoded probability for all times when a rat was at a given position was calculated. To determine the decoding error, the decoded position was defined as the position with maximal decoded probability for each time bin. The decoding error was then defined as the difference between a rat’s actual position and the corresponding decoded position. The cumulative error distributions for each day were then determined (Figure 4A-B). For a day of recordings to be included in theta sequence event and replay event analysis, its cumulative error distribution had to reach 50% at error values less than 20 degrees (Davidson et al., 2009; Hwaun & Colgin, 2019). Days that reached this criterion are shown in Figure 4A-B. The overall error distributions for rats with days that met this criterion are shown in Figure 4C.

**Figure 4.**
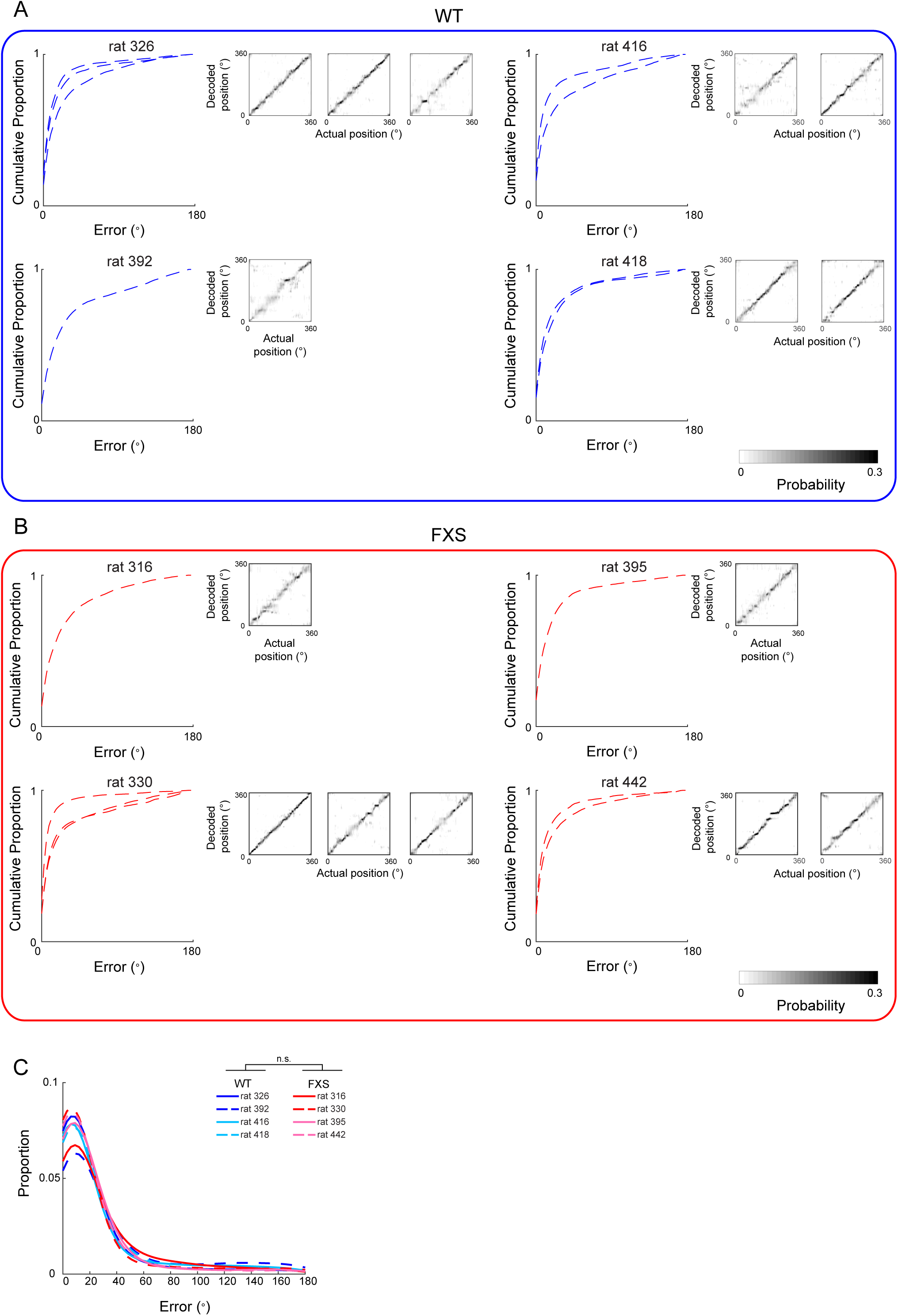
Decoding accuracy in WT and FXS rats. (A-B) Cumulative decoding error and confusion matrices are shown for decoded place cell populations from WT (A) and FXS (B) rats. Individual lines on cumulative error plots represent individual days. Insets show confusion matrices from each day for each rat. Only days that met the decoding criteria are shown. (C) There was no difference in decoding error values between WT and FXS rats. Each line represents the distribution of decoding error values from one rat.

### Detection and analysis of theta sequences

Theta sequences were detected only on days that reached the decoding criterion (see “*Decoding accuracy analysis”* section above). Theta sequences were detected and characterized using methods similar to our previously published procedures (Zheng et al., 2016). LFP signals were band-pass filtered in the theta band (6-10 Hz), and individual theta cycles were cut at the theta phase with the lowest number of spikes. Bayesian decoding was then performed on theta cycles in which at least 3 place cells fired and when the rat was traveling at a speed of 5 cm/s or greater (see “*Bayesian decoding analysis”* section above). Bayesian decoding was performed across partially overlapping 40 ms windows advanced by a 10 ms step (Figure 5A-B). If spikes did not occur throughout the entirely of the theta cycle, contiguity was considered to be broken when there were two consecutive time bins (i.e., 50 ms) without spikes. Sequence property and significance analyses were then performed on the longest set of contiguous time bins (Figure 5C-F). The t-span of a sequence was defined as the temporal duration of these time bins. To determine the slope and x-span of the sequence event, we calculated the center of mass of the posterior probability distribution *P(x | n)* from the Bayesian decoding for each time bin. A circular-linear regression line was fit to these positions in order to determine the slope. The x-span was defined as the distance between the first and last positions of the regression line. To determine if a theta sequence was significant, we compared the r^2^ value from this circular-linear regression to a shuffled distribution. This distribution was obtained by circularly shuffling each time bin of the weighted probability distribution (i.e., *P(x | n)* from the Bayesian decoding analysis) a random distance 1000 times and then obtaining an r^2^ value for each shuffled distribution. The r^2^ value from the theta sequence had to exceed 95% of the shuffled values in order to be considered significant. Additionally, to ensure the regression provided an accurate representation of the true decoded probability distribution for the event, at least 60% of the total posterior probability needed to be within 20 degrees of the fitted circular-linear regression line (Zheng et al., 2021). In addition, the minimum distance between the fitted trajectory and the actual position of the rat had to be less than 20 degrees (Zheng et al., 2021). The number of theta sequences from each rat is shown in Table 3.

**Figure 5.**
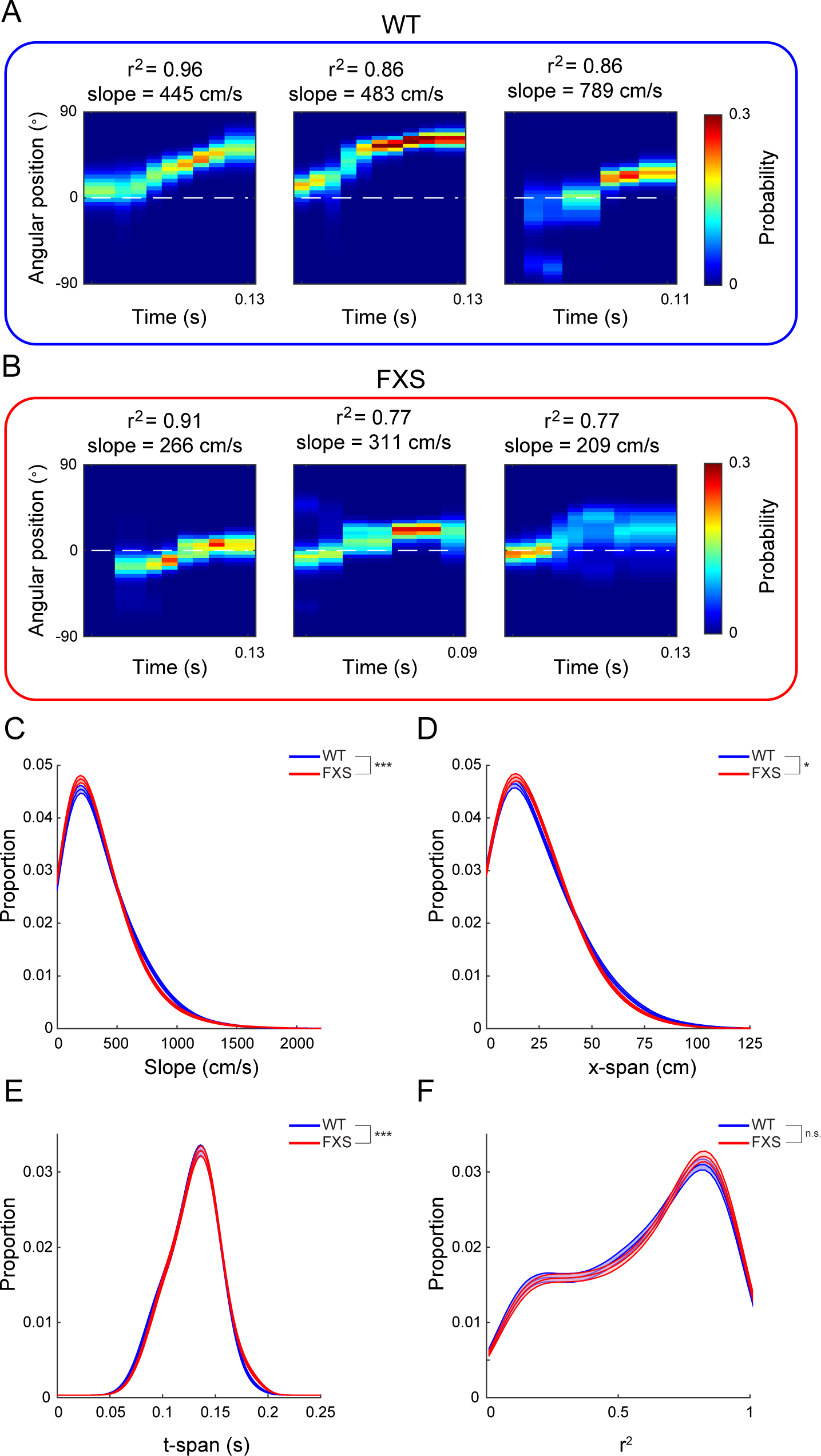
Theta sequence events coded paths that were less temporally compressed and shorter in FXS rats than in WT rats. (A-B) Example theta sequence events are shown for WT (A) and FXS (B) rats. Position (on the y-axis) is shown relative to the mean of the rat’s actual position during the theta sequence event (indicated by white dashed line). The associated r^2^ and slope values are shown for each event. (C) Theta sequences’ slopes were lower in FXS rats than WT rats, indicating that theta sequences were less temporally compressed in FXS rats. (D) The x-span values of theta sequences were lower in FXS rats than in WT rats, indicating that theta sequences represented relatively short spatial paths in FXS rats. (E) The duration (t-span) of theta sequences was higher in FXS rats than in WT rats. (F) There was no difference in r^2^ values between FXS and WT rats, indicating that theta sequences represented coherent paths through an environment in FXS rats despite the reduced temporal compression of representations of these paths. (C-F) The distributions for theta sequence properties are shown with shaded areas representing 95% confidence intervals of the distributions for each genotype.

**Table 3.**
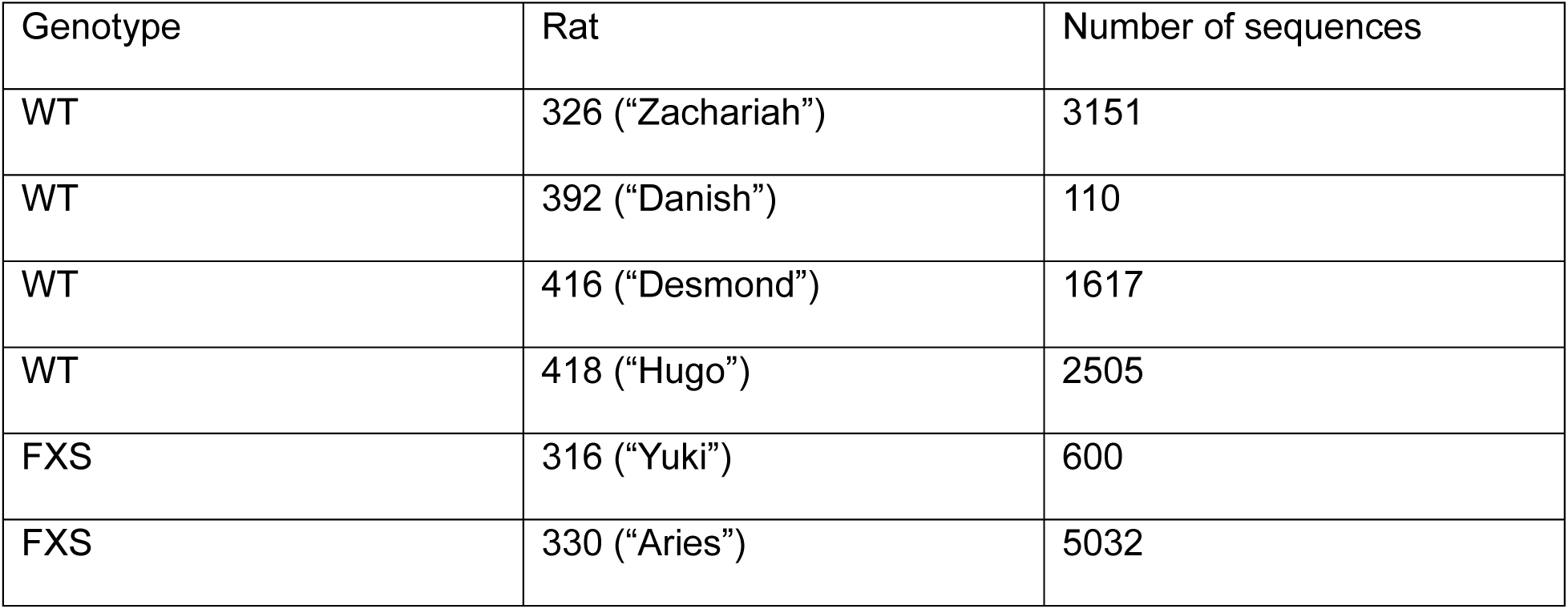
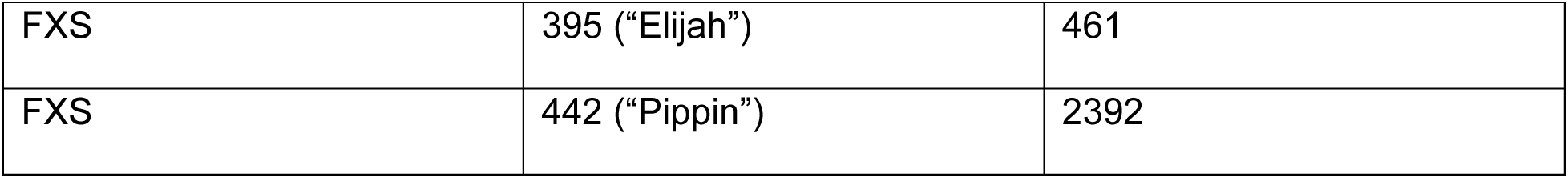
Total number of theta sequences recorded from each rat.

### Detection and characterization of replay events

Replay events were detected only on days that reached the decoding criterion (see “*Decoding accuracy analysis”* section above). Replay events were detected using methods similar to previously published procedures (Hwaun & Colgin, 2019; Pfeiffer & Foster, 2013). Replay events were detected while the rat rested off of the circular track in a towel-lined flowerpot (see “*Behavior”* section above). To detect candidate events, a histogram of population firing rates was constructed using all cells that were classified as active during the track running sessions (see “*Place cell analyses”* section above). This histogram was then smoothed with a Gaussian kernel (standard deviation = 10 ms). Candidate events were detected when the population firing rate exceeded 3 standard deviations above the mean population firing rate and were bounded by crossing of the mean. Events within 40 ms of each other were combined. Start and end times were then adjusted inward so that the first and last time bins of a candidate event each contained at least one spike. To be included for further analysis, an event had to be between 50 and 2000 ms in duration, and at least 5 cells had to fire during an event.

For each candidate event, Bayesian decoding was performed (see “*Bayesian decoding analysis*” section above) across partially overlapping 20 ms windows advanced by a 10 ms step (Figure 6A-B). To assess how well a posterior probability distribution represented an actual trajectory on the circular track, circular-linear regression was performed with the decoded position as the circular variable and time within the event as the linear variable. The decoded position for each time bin was defined as the center of mass of each time bin, which was determined by taking the circular mean of positions weighted by their associated posterior probability values. The r^2^ value of the regression line was used as an assessment of replay fidelity (Figure 6C), as in previous studies (Davidson et al., 2009; Hwaun & Colgin, 2019; Karlsson & Frank, 2009). To be classified as a significant replay event, the r^2^ value of an event’s regression line had to be at least 0.5 (as in Hwaun & Colgin, 2019), and the decoded positions between adjacent time bins (i.e., the “jump” distance between adjacent time bins) could not exceed 25% of the length of the circular track (as in Berners-Lee et al., 2022). Only replay events that occurred after the first track running session of the day (i.e., in rest sessions 2-5) were included for further analysis. To quantify the temporal compression of a replay event, we estimated the slope of the circular-linear regression line (Figure 6E). We calculated the absolute value of the slope to include both forward and reverse replay events. The path distance represented within a replay event was calculated as the difference between the decoded positions of the first and last time bins of a replay event (Figure 6F).

**Figure 6.**
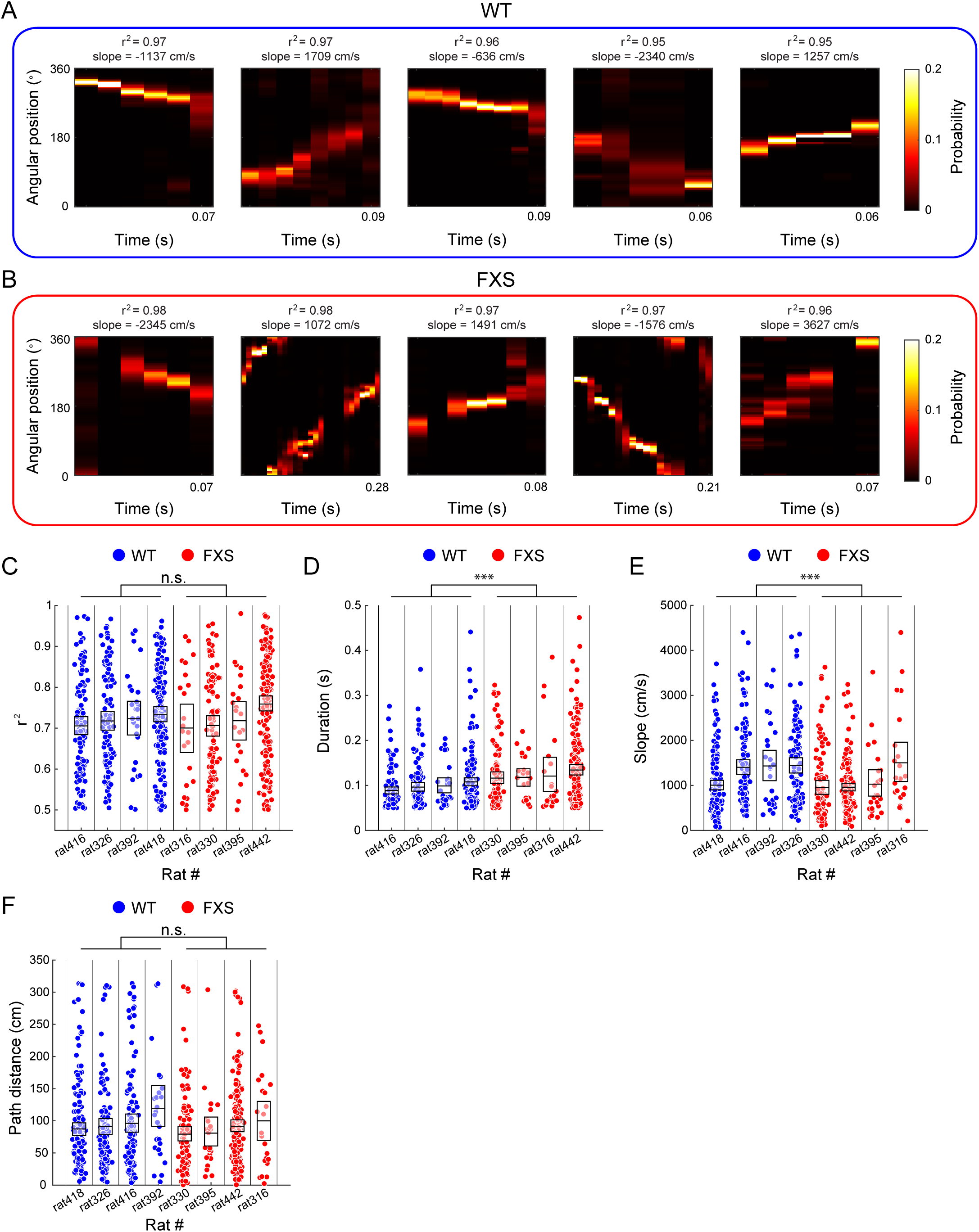
Replay events were less temporally compressed in FXS rats than in WT rats. (A-B) Example replay events are shown for WT (A) and FXS (B) rats. The r^2^ value and slope of each replay event is shown above each plot. (C) Replay events’ r^2^ values did not differ between WT and FXS rats. (D) Replay event durations were longer in FXS rats than in WT rats. (E) The slopes of the regression lines fit to posterior probability distributions of replay events were lower in FXS rats than in WT rats. (F) Path distances of replayed trajectories did not differ between WT and FXS rats. (C-F) Each dot represents a measure for one replay event. Boxes represent 95% confidence intervals for the mean values from each rat.

### Replay event place cell analyses

To investigate the firing of place cells during replay events, we binned the firing rates of CA1 place cells in 1 ms bins around times of replay event onset for each unit (as in Hwaun & Colgin, 2019). We only included replay events in which a unit participated (as in Boone et al., 2018). This allowed us to examine differences in place cell firing patterns during replay events while controlling for differences in the number of replay events in which a unit participates. We smoothed each firing rate vector with a Gaussian kernel (standard deviation = 10 ms). We averaged across all replay events to obtain a binned mean firing rate vector for each unit (Figure 7A). Additionally, we determined the average firing rate for each unit during all replay events in which a unit participated (Figure 7B) and the average number of spikes per event that each unit fired (Figure 7C) (as in Boone et al., 2018).

**Figure 7.**
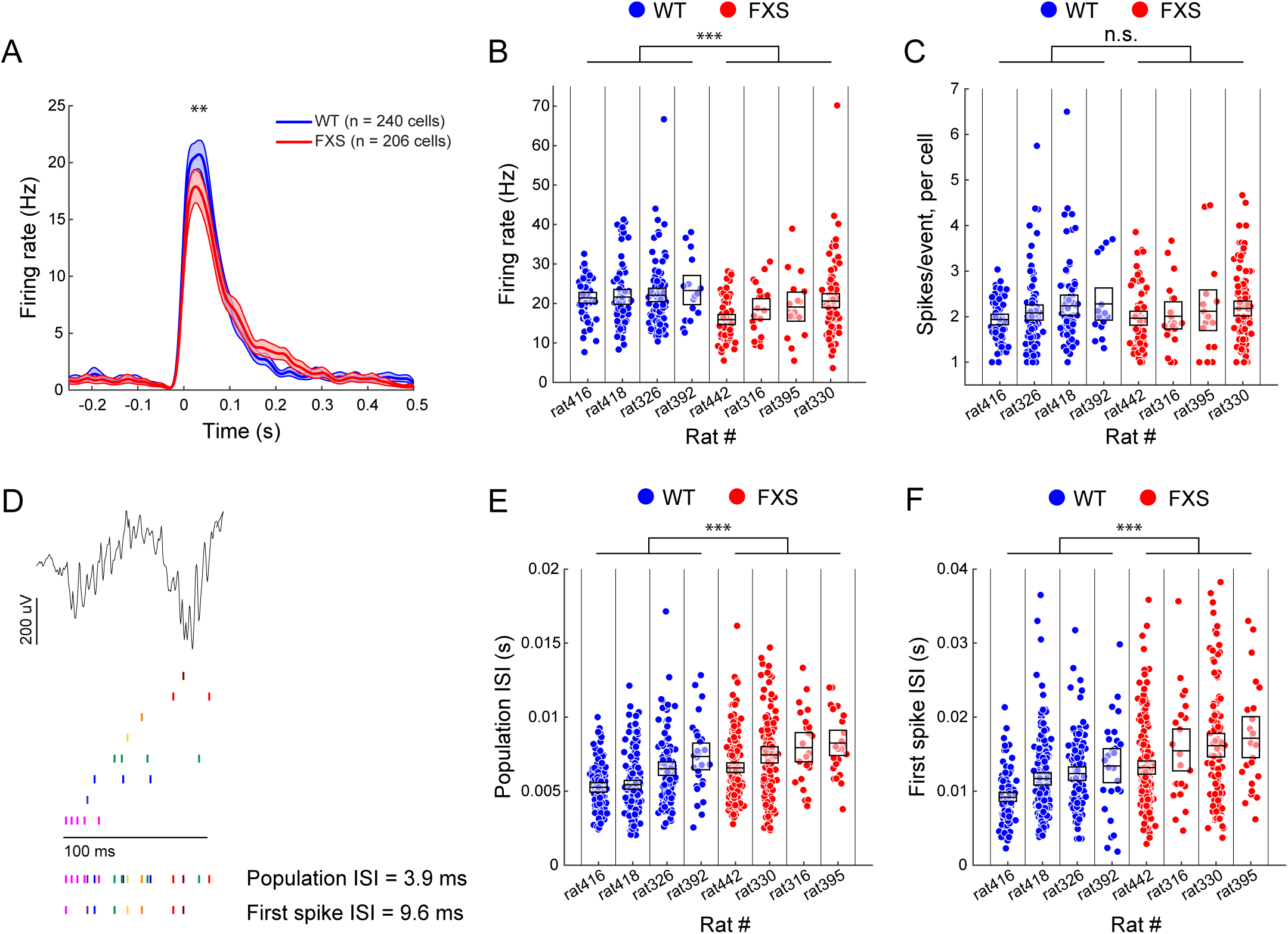
Place cells fired more slowly during replay events in FXS rats than in WT rats. (A) Place cells from WT rats reached a higher peak firing rate after replay event onset than place cells from FXS rats. Shaded areas represent 95% confidence intervals of the binned firing rate distributions. (B) Place cell firing rates during replay events were higher in WT rats than in FXS rats. (C) The number of spikes a place cell fired during replay events did not differ between WT and FXS rats. (D) Schematic illustrating how population inter-spike interval (ISI) and first spike ISI were calculated. An LFP recording during an example replay event from a WT rat is shown (top). A raster plot shows spiking activity of 8 place cells that participated in the replay event, with spikes from different cells represented by different colored tick marks (middle). The bottom two rows show spikes included when calculating the average ISI for all spikes (population ISI) and only the first spike from each cell (first spike ISI). (E) Population ISIs were higher in FXS rats than in WT rats. (F) First spike ISIs were higher in FXS rats than in WT rats. (B-C) Each dot represents a measure for an individual place cell. Boxes represent 95% confidence intervals for the mean values for each rat. (E-F) Each dot represents one replay event. Boxes represent 95% confidence intervals for the mean values for each rat.

To determine whether the spike timing of place cells during replay events was impaired in FXS rats, we calculated the interval between spikes in a replay event in two ways (Figure 7D). For the first method, the “population inter-spike interval”, we calculated the average interval between consecutive spikes from all cells in a replay event for each replay event (Figure 7E). In the second method, the “first spike inter-spike interval”, we only considered the first spike that each cell fired in a replay event and then computed the average interval between consecutive times of first spikes for each replay event (Figure 7F).

### Replay event LFP analyses

The LFP from the tetrode with the most place cells on a given day was used for each day’s analysis of power associated with replay events. The time-varying power across frequencies around the time of replay event onset was computed using a wavelet transform (Tallon-Baudry et al., 1997) as previously described (Mably et al., 2017). Time-varying power around replay event onset was calculated in 1 Hz steps from 2-250 Hz and averaged across all replay events within a day. Power was z-scored within each frequency band. A time-varying power vector for each frequency was created for each rat by averaging across recording days. The plots presented in Figure 8A-B represent the average time-frequency representations of power associated with replay events across all rats for each genotype. The peak ripple frequency for a replay event (Figure 8C) was defined as the frequency with the highest power in the ripple band, 150-250 Hz. To calculate an overall estimate of slow gamma power associated with a replay event (Figure 8D), we averaged power estimates across the slow gamma band of frequencies (25-55 Hz) and across time for each replay event.

**Figure 8.**
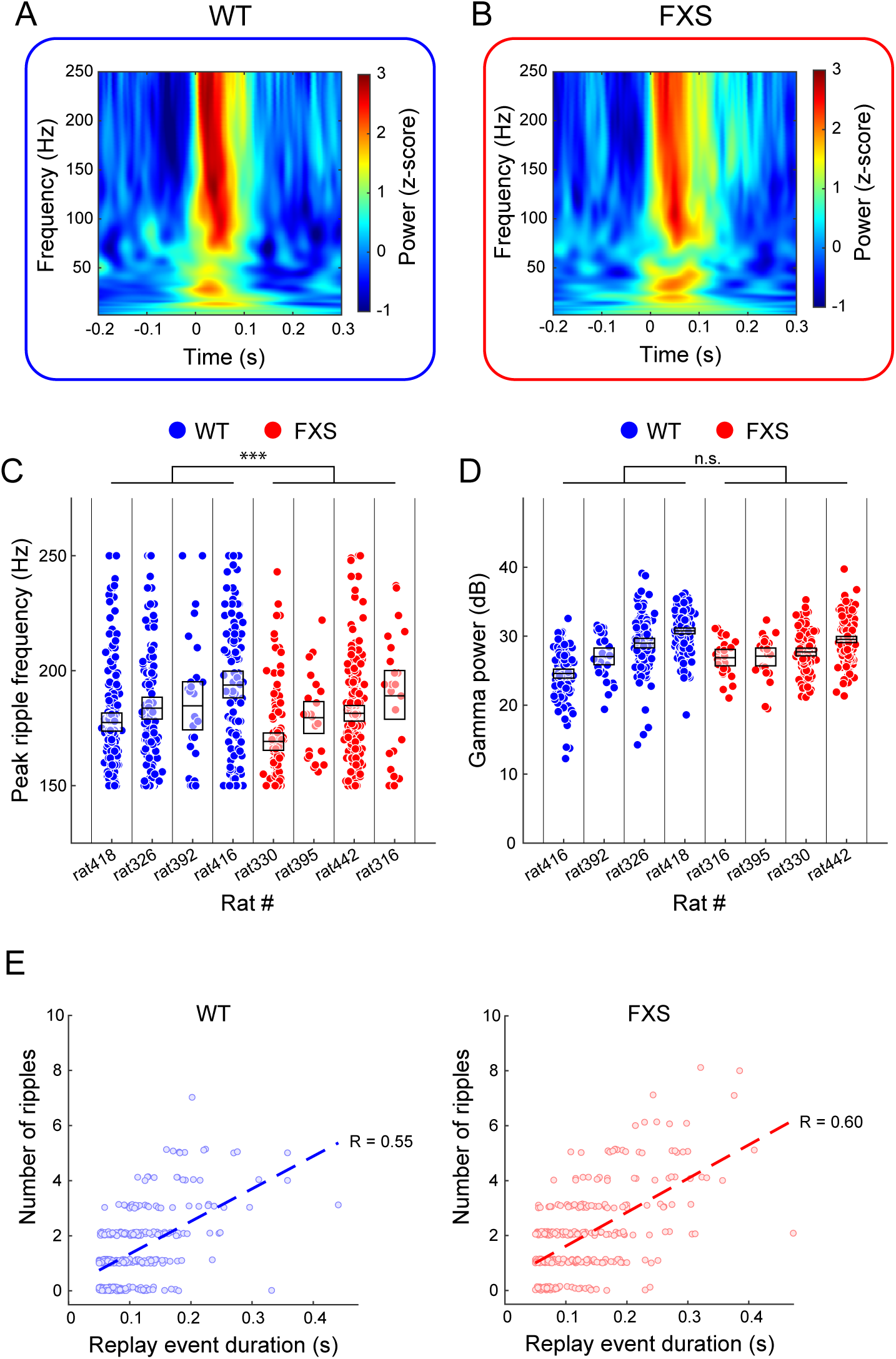
Peak ripple frequency was lower in FXS rats than in WT rats. (A-B) Time-frequency representations of power during replay events in WT (A) and FXS (B) rats. (C) Peak ripple frequency was lower in FXS rats than in WT rats. (D) Slow gamma power during replay events (25-55 Hz) did not differ between WT and FXS rats. (C-D) Each dot represents a measure for one replay event. Boxes represent 95% confidence intervals for the mean values for each rat. (E) The number of ripples that co-occurred with replay events of a given duration did not differ between WT (left) and FXS (right) rats. Each dot represents a measure for one replay event. The dashed line represents the regression line fit to the distribution of points for each genotype.

To assess the relationship between the duration of replay events and number of sharp wave-ripples (Figure 8E), we used methods adapted from a previous study (Davidson et al., 2009). The LFP from the tetrode with the most place cells from a given day was bandpass filtered from 150-250 Hz, and a Hilbert transform was then applied to the filtered signal. The absolute value of the Hilbert transformed signal was then smoothed with a Gaussian kernel (standard deviation = 8 ms) and z-scored. Ripples were detected at times when the z-score was greater than or equal to 3. The number of ripples is displayed with a small amount of random jitter on the y-axis for visualization (Figure 8E) (as in Davidson et al., 2009).

### Data visualization

Whenever possible, data is shown for each animal individually. In Figures 2, 3, 6, 7, and 8, each data point represents one measure (i.e., for an individual place cell or replay event). Boxes show 95% confidence intervals of the mean values within each rat. Confidence intervals were calculated with 1000 bootstrapped samples. WT rats are shown with blue data points, and FXS rats are shown with red data points. In each plot for each genotype, data from individual rats were presented in an order corresponding to increasing mean values for ease of comparison across genotypes. In Figures 4 and 5, distributions are shown due to the large number of data points for each genotype. In Figure 5, shaded areas represent 95% confidence intervals calculated with 1000 bootstrapped samples.

### Experimental design and statistical analyses

Whenever possible, experimenters were blinded to genotype during data acquisition and analyses. However, due to temporarily limited availability of FXS rats from the supplier, experiments on two cohorts of rats were performed unblinded (rats 416, 418, 442, and 445).

MATLAB (Mathworks) scripts were custom written for the analyses in this study, based on algorithms that have been used in prior studies, as described above. All statistical tests were performed using SPSS Statistics (Version 29.0, IBM) unless otherwise stated. To compare behavior, place cell properties, decoding errors, and replay properties across genotypes, we used a generalized linear mixed model design (as in Robson et al., 2024). Individual rats were subjects, and genotype was included as a fixed factor. When comparing behavior metrics, sessions were nested within days as repeated measures. When applicable, place cells were nested within rats (analyses in Figures 2, 3, and 7A-C). When comparing different session pairs to assess place cell firing rate map stability (Figure 2A-B), session pair was included as a repeated measure. To compare decoding errors between genotypes (Figure 4C), the error calculated for each time bin in which a rat was moving faster than 5 cm/s (see “*Decoding accuracy analysis”* section above) was nested within sessions, and sessions were nested within days as repeated measures. For analyses of replay event properties (Figures 6, 7E-F, and 8C-D), individual replay events were nested within the rest session in which they occurred, and rest sessions were nested within a day as repeated measures (as in Boone et al., 2018).

To compare theta sequence properties between genotypes (Figure 5C-F), we used the non-parametric Wilcoxon rank-sum test (*ranksum* function in MATLAB).

To account for potential differences in decoding accuracy across rats when analyzing theta sequence slope and replay event slope, we used a generalized linear model (*fitglm* function in MATLAB) with a log-link function. The model included genotype and decoding errors for each session as predictors of slope values to determine if decoding accuracy significantly affected the results (similar to analysis of effects of decoding accuracy in Hwaun & Colgin, 2019).

To assess whether the correlation between replay duration and number of sharp wave-ripples differed across genotypes (Figure 8E), we used multiple linear regression. Genotype, replay event duration, and the interaction between genotype and replay event duration were included as predictors.

### Code and data accessibility

Analysis code is available on GitHub (https://github.com/mmdonahue/FMR1CircTrack_PublicShare). Data will be made available upon reasonable request.

## Results

### Hippocampal place cells showed similar activity in FXS and WT rats

To determine how place cell coding of spatial locations is affected in a rat model of FXS, we recorded from the CA1 pyramidal cell layer of the hippocampus in adult FXS and WT rats. We recorded place cell activity while rats ran unidirectionally on a circular track in a familiar room for four ten-minute sessions per day (Figure 1). WT and FXS rats ran at similar speeds on the circular track (generalized linear mixed model, WT: estimated mean = 34 cm/s, estimated 95% CI = 28-39 cm/s; FXS: estimated mean = 27 cm/s, estimated 95% CI = 22-33 cm/s; no significant main effect of genotype (F(1,98) = 2.5, p = 0.116)), However, FXS rats completed fewer laps per session on average (generalized linear mixed model, WT: estimated mean = 17 laps, estimated 95% CI = 13-22 laps; FXS: estimated mean = 10 laps, estimated 95% CI = 5-15 laps; significant main effect of genotype (F(1,98) = 4.4, p = 0.039), suggesting that FXS rats paused on the track more than WT rats.

Previous work in a different FXS rat model (i.e., Long Evans *Fmr1* knockout rats) found no difference in spatial stability of CA1 place cells when rats explored an initially novel environment over two days (Asiminas et al., 2022). However, work from FXS mice reported impaired short-term stability in CA1 place cells (Arbab, Pennartz, et al., 2018). To examine place cell stability in the present FXS rat model, we computed spatial correlation coefficients between pairs of track running sessions for each place cell. Spatial correlations were lower in FXS rats than in WT rats, suggesting impaired place field stability in FXS rats (Figure 2A; generalized linear mixed model, no significant interaction between genotype and session pair (F(3,2036) = 0.399, p = 0.754), significant main effect of genotype (F(1,2036) = 4.596, p = 0.032)). Further, place cell firing rates changed more between sessions in FXS rats than in WT rats (Figure 2B; generalized linear mixed model, no significant interaction between genotype and session pair (F(3,2036) = 0.615, p = 0.605), significant main effect of genotype (F(1,2036) = 69.542, p < 0.001)). The seeming discrepancy between our results and the relatively stable place fields observed in another rat model of FXS (Asiminas et al., 2022) may be due to differences in testing environments or rat strains.

The previous study of CA1 place cell activity in a different FXS rat model than the one used in the present study showed no impairments in place cells’ spatial information, mean firing rates, or place field size (Asiminas et al., 2022). Here, we also found no differences between FXS and WT rats in CA1 place cells’ spatial information (Figure 2C; generalized linear mixed model, no significant main effect of genotype (F(1,436) = 3.674, p = 0.056)), peak firing rates (Figure 2D; generalized linear mixed model, no significant main effect of genotype (F(1,458) = 0.316, p = 0.574)), and place field size (Figure 2E; generalized linear mixed model, no significant main effect of genotype (F(1,458) < 0.001, p = 0.995)). This indicates that CA1 place cells in FXS rats can represent different locations within a session in a familiar environment as well as CA1 place cells in WT control rats.

### Theta modulation of place cells was normal in FXS rats

During active exploration, the activity of place cells is coordinated by the ∼6-10 Hz theta rhythm in the hippocampus. Theta phase precession refers to a place cell firing pattern in which successive place cell spikes occur at progressively earlier phases of the theta cycle as a rat traverses the cell’s place field (O’Keefe & Recce, 1993). Manipulations that reduce theta phase precession (Robbe & Buzsáki, 2009) and disrupt the precise spike timing of neurons during theta sequences (Petersen & Buzsáki, 2020) reduce spatial memory performance, raising the possibility that abnormal theta phase precession contributes to spatial memory deficits that have been reported in FXS models (Asiminas et al., 2019; Bakker et al., 1994; Mineur et al., 2002; Tian et al., 2017; Till et al., 2015; Van Dam et al., 2000). We examined whether CA1 place cells exhibit abnormal phase precession in FXS rats (Figure 3). We found no difference between FXS and WT rats in the relationship between theta phase and relative position in the place field (Figure 3C; generalized linear mixed model, no significant main effect of genotype (F(1,423) = 0.795, p = 0.373)), suggesting that theta coordination of spiking is normal in FXS rats at the level of individual cells.

### Theta sequences spanned shorter distances in FXS rats

Other work has led to the hypothesis that individual place cells are largely normal in rodent models of FXS while coordinated activity of large populations of place cells is impaired (Talbot et al., 2018). Studies in FXS mice have suggested that hippocampal networks are hypersynchronized in FXS (Arbab, Battaglia, et al., 2018; Talbot et al., 2018). FXS mice also showed abnormally low coordination between place cells with overlapping place fields (Talbot et al., 2018). Together, these prior results suggest that coordinated theta sequences of place cells may be impaired in FXS rats. To assess theta sequences in FXS rats, we used a Bayesian decoding approach (K. Zhang et al., 1998). Specifically, we applied Bayesian decoding to spiking activity from place cell populations during active running sessions to reconstruct positions on the circular track represented by place cell populations. We first determined how accurately we were able to reconstruct the rat’s positions during running using the place cell spiking activity. Most rats’ recordings (WT rats 326, 392, 416, and 418 and FXS rats 316, 330, 395, and 442) surpassed our criteria for sufficient decoding accuracy (Figure 4A-B; see also “*Decoding accuracy analysis*” section in Materials and Methods). Also, decoding errors were not significantly different between genotypes (Figure 4C; generalized linear mixed model, no significant main effect of genotype (F(1,573941) = 0.17, p = 0.678)).

For the animals that surpassed decoding accuracy criteria, we used Bayesian decoding to identify trajectories represented by place cell populations within individual theta cycles as WT and FXS rats ran laps unidirectionally on a circular track (Figure 5A-B; for the number of theta sequences per rat, see Table 3). We then performed circular-linear regression to fit a regression line to the spatial path represented within these theta cycles. This regression line allowed us to quantify the temporal compression (slope, in cm/s), distance (“x-span”, in cm), and duration (“t-span”, in s) of each trajectory. We found that the slopes of theta sequences were lower in FXS rats (Figure 5C; Wilcoxon rank-sum test, significant effect of genotype (Z = 3.7, p < 0.001)). To account for potential differences in decoding accuracy across different rats and days, we used a generalized linear model with a log-link function to predict slope values from genotype and the decoding errors for each session. Including genotype in the model increased prediction accuracy while including decoding errors had no significant effect on the model (genotype, β = 0.2, p = 0.007; decoding accuracy, β = 0.08, p = 0.32), suggesting that the difference in the temporal compression of theta sequences was not due to differences in decoding accuracy between genotypes. The difference in slope values reflected both a decrease in the distances of paths represented during theta sequences (Figure 5D; Wilcoxon rank-sum test, significant effect of genotype (Z = 2.2, p = 0.029)) and an increase in the duration of events (Figure 5E; Wilcoxon rank-sum test, significant effect of genotype (Z = 3.6, p < 0.001)). We also calculated the r^2^ value of the regression line fit to decoded sequence representations in order to assess the extent to which sequences represented coherent trajectories. We found no significant difference in the r^2^ values of the circular-linear regression lines for sequences from WT and FXS rats (Figure 5F; Wilcoxon rank-sum test, no significant effect of genotype (Z = 1.9, p = 0.062)).

These results suggest that theta sequences represented spatial trajectories in FXS rats. However, theta sequences represented shorter trajectories, and representations were less temporally compressed, in FXS rats compared to WT rats.

### Replay events were less temporally compressed in FXS rats

Replay events co-occur with sharp wave-ripples in the LFP of the hippocampus during NREM sleep or waking rest (Davidson et al., 2009; Kudrimoti et al., 1999; Lee & Wilson, 2002; Nádasdy et al., 1999). Previous work has shown that sharp wave-ripples during NREM sleep are abnormally long in duration in FXS mice (Boone et al., 2018). To assess replay events in FXS rats, we examined the firing of place cell sequences while rats rested quietly in a location off the circular track after running. We used Bayesian decoding to reconstruct positions on the circular track represented by CA1 place cell populations during replay events in WT and FXS rats (Figure 6A-B; for the number of replay events recorded from each rat, see Table 4). We quantified replay fidelity, duration, and temporal compression of each replay event using protocols similar to previously published procedures (Davidson et al., 2009; Hwaun & Colgin, 2019; Karlsson & Frank, 2009). Specifically, we first applied circular-linear regression analysis to the posterior probability distributions resulting from Bayesian decoding of place cell spikes during replay events. We then assessed associated r^2^ values as a measure of replay fidelity, and the slopes of the fitted lines were used to estimate the temporal compression of replay events. There was no difference in replay fidelity between WT and FXS rats (Figure 6C; generalized linear mixed model, no significant effect of genotype (F(1,751) = 2.185, p = 0.140)). However, the duration of replay events was longer in FXS rats than in WT rats (Figure 6D; generalized linear mixed model, significant effect of genotype (F(1,751) = 33.332, p < 0.001)).

**Table 4.** Total number of replay events from each rat.

| Genotype | Rat | Number of replay events |
| --- | --- | --- |
| WT | 326 ("Zachariah") | 114 |
| WT | 392 ("Danish") | 28 |
| WT | 416 ("Desmond") | 116 |
| WT | 418 ("Hugo") | 166 |
| FXS | 316 ("Yuki") | 23 |
| FXS | 330 ("Aries") | 110 |
| FXS | 395 ("Elijah") | 24 |
| FXS | 442 ("Pippin") | 172 |

Further, the representations during replay events were less temporally compressed, indicating slower transitions across representations of successive locations, in FXS rats than in WT rats (Figure 6E; generalized linear mixed model, significant effect of genotype (F(1,751) = 19.340, p < 0.001)). To account for potential differences in decoding accuracy across different rats and days, we used a generalized linear model with a log-link function to predict slope values from genotype and the decoding errors for each session. Including genotype in the model increased prediction accuracy while including decoding errors had no significant effect on the model (genotype, β = 0.25, p < 0.001; decoding accuracy, β = 0.06, p = 0.82), suggesting that the difference in temporal compression of replay events was not due to different decoding accuracy between genotypes. The low temporal compression in FXS rats did not indicate that replay events represented unusually long paths in FXS rats, however, because path lengths of replay sequences were similar in FXS and WT rats (Figure 6F; generalized linear mixed model, no significant effect of genotype (F(1,751) = 1.3, p = 0.253). Instead, this pattern of results suggests that replay events in FXS rats represented trajectories of similar lengths and comparable fidelity as in WT rats but that replay of representations of trajectories occurred more slowly in FXS rats than in WT rats.

### Place cells fired more slowly during replay events in FXS rats

In addition to differences in duration, Boone and colleagues (Boone et al., 2018) observed abnormal place cell firing during sharp wave-ripples in a mouse model of FXS. They found that place cells had lower in-event firing rates during sharp wave-ripples recorded during sleep, although individual cells fired the same number of spikes per event (Boone et al., 2018). Abnormally slow firing of place cells during replay events in FXS rats may underlie the reduced temporal compression observed in our data. Indeed, we found that CA1 place cells had lower peak firing rates after replay event onset in FXS rats than in WT rats (Figure 7A; generalized linear mixed model, significant effect of genotype on peak firing rate after event onset (F(1,444) = 8.877, p = 0.003)). In-event firing rates were also lower in FXS rats than in WT rats (Figure 7B; generalized linear mixed model, significant effect of genotype (F(1,444) = 23.140, p < 0.001)), while the number of spikes each cell fired during a replay event did not differ between FXS and WT rats (Figure 7C; generalized linear mixed model, no significant effect of genotype (F(1,444) = 1.020, p = 0.313)) likely because of replay events’ relatively long durations in FXS rats (Figure 6D). The low replay-associated firing rates in individual place cells in FXS rats suggest that dynamics of CA1 place cell sequence firing may occur more slowly during replay events in FXS rats than in WT rats. To test this hypothesis, we calculated the distribution of intervals between successive spikes in a replay event in two ways (Figure 7D). First, we considered all spikes that occurred across all cells during a replay event and found that these population inter-spike intervals were longer in FXS rats than in WT rats (Figure 7E; generalized linear mixed model, significant effect of genotype (F(1,751) = 58.337, p < 0.001)). Next, we only considered the first spike that each cell in a sequence fired in a replay event (Figure 7F). These first spike inter-spike intervals were also longer in FXS rats than in WT rats (generalized linear mixed model, significant effect of genotype (F(1,751) = 56.843, p < 0.001)).

### Properties of ripples and slow gamma rhythms during replay events in FXS rats

We further examined properties of oscillatory patterns in the LFP, specifically sharp wave-ripples and slow gamma oscillations, that normally co-occur with replay events in WT rats (Davidson et al., 2009; Kudrimoti et al., 1999; Lee & Wilson, 2002; Nádasdy et al., 1999; Carr et al., 2012; Bieri et al., 2014). In both genotypes, we saw increases in power in the ripple and slow gamma bands at event onset (Figure 8A-B), suggesting that ripples and slow gamma rhythms co-occur with replay events in FXS rats as in WT rats. However, the peak ripple frequency during replay events was lower in FXS rats than in WT rats (Figure 8C; generalized linear mixed model, significant effect of genotype (F(1,751) = 21.168, p < 0.001)), consistent with results from FXS mice showing lower peak ripple frequency during sharp wave-ripples during sleep (Boone et al., 2018). Although the frequency of ripples was lower in FXS rats, the number of sharp wave-ripples that co-occurred with a replay event of a given duration was similar in FXS and WT rats (Figure 8E; significant multiple linear regression (F(3,749) = 145.149, p < 0.001); no significant interaction between genotype and replay event duration (t = 0.369, p = 0.712)), likely due to the longer duration of replay events in FXS rats than in WT rats (Figure 6D).

Previous work has shown increased slow gamma power in CA1 during sleep-associated sharp wave-ripples in a FXS mouse model (Boone et al., 2018). Here, we found no difference in slow gamma power during rest-associated replay events in WT and FXS rats (Figure 8D; generalized linear mixed model, no significant effect of genotype (F(1,751) = 0.095, p = 0.758)). Replay events during sleep and waking rest may have different functions and characteristics (Roumis & Frank, 2015; Tang et al., 2017) and may involve input from different upstream structures to CA1 (Yamamoto & Tonegawa, 2017). Thus, differences in rest and sleep states may underlie the seemingly discrepant results for replay-associated slow gamma in FXS rat and mouse models.

## Discussion

To our knowledge, this is the first study that examines spatial representations coded by large populations of hippocampal place cells in a rodent model of FXS. Here, we present data showing that coding of spatial trajectories by coordinated place cell populations is impaired in FXS rats. Specifically, we found that theta sequences coded less temporally compressed representations and represented shorter path distances in FXS rats, while theta phase precession in individual place cells was normal. Further, we found that place cell sequences fired abnormally slowly during hippocampal replay events in FXS rats. This slow place cell firing was associated with reduced temporal compression of representations of track trajectories during replay in FXS rats. Together, these results suggest that coordinated place cell sequences code spatial trajectories abnormally during both active running and during subsequent rest in a rat model of FXS.

Previous studies examining the activity of place cells during active exploration in FXS rodents have yielded mixed results. Studies from FXS mice and rats have shown no difference in spatial information in CA1 place cells (Arbab, Pennartz, et al., 2018; Asiminas et al., 2022; Talbot et al., 2018), although one study has shown abnormally large place field sizes and excessive out-of-field firing in FXS mice (Arbab, Pennartz, et al., 2018). Discrepancies between these results and the present findings may result from different experimental paradigms, including different testing arenas, experimental time courses, and the degree of environmental novelty. Regarding the latter point, work from FXS rats has shown that experience-dependent reductions in firing rates and sharpening of spatial tuning of place cells are lacking in FXS rats after introduction to a novel environment (Asiminas et al., 2022). These results suggest that the initial formation of a spatial memory may be stunted in FXS rats. We did not observe such differences in our study, likely due to our use of a testing environment with which rats had already been familiarized.

Work from FXS mice has shown normal theta modulation in CA1 place cells but reduced correlated firing of pairs of place cells (Talbot et al., 2018). This suggested that coordinated population coding would be impaired in FXS rodents. Consistent with this result, we show that theta-coordinated sequences of place cells during active exploration represented shorter trajectories in FXS rats, while theta phase precession in individual place cells remained normal. While theta phase precession and theta sequences appear to be related phenomena, it is possible to observe the absence of theta sequences in populations of place cells that individually show intact phase precession (Feng et al., 2015; Middleton & McHugh, 2016).

Computational models have suggested that alterations in the coupling between place cells and local CA1 interneurons may disrupt the temporal compression of theta sequences while leaving phase precession intact (Chadwick et al., 2016). Interneurons are less modulated by theta in FXS mice (Talbot et al., 2018), and CA1 interneurons and pyramidal cells are less correlated in FXS mice (Arbab, Battaglia, et al., 2018). This interneuron-pyramidal cell dysfunction may result in hyper-synchronization between CA1 pyramidal neurons and thereby cause place cells that represent multiple locations along a trajectory to fire more synchronously in FXS rats. This could flatten slopes of theta sequences and shorten distances of represented trajectories.

Disrupting the input from CA3 to CA1 may also affect the path length of theta sequences (Chadwick et al., 2016; Middleton & McHugh 2016). *In vitro* work examining the Schaffer collateral pathway from CA3 to CA1 in FXS has yielded mixed results. Work from FXS mice has shown an abnormally low induction threshold for long-term potentiation (LTP) in FXS mice when pre– and post-synaptic neurons are simultaneously activated (Routh et al., 2013). However, other work from FXS mice (Lauterborn et al., 2007) and rats (Tian et al., 2017) has reported reduced Schaffer collateral LTP. FXS mouse models have also shown increased metabotropic glutamate receptor (mGluR) long-term depression (LTD) (Huber et al., 2002; Iliff et al., 2013; J. Zhang et al., 2009), suggesting that hippocampal synapses may be relatively weak in rodent models of FXS. Such a reduction in synaptic input from CA3 to CA1 may affect the temporal compression of theta sequences in FXS rats, resulting in representation of relatively short paths.

Our work shows that CA1 place cells fired more slowly in FXS rats than in WT rats during replay events that occurred during waking rest. This finding is consistent with abnormally slow firing of pyramidal cells during sharp wave-ripples in non-REM sleep reported in FXS mice (Boone et al., 2018). *In vitro* work from FXS rats has also shown lower frequency multi-unit activity during sharp wave-ripples specifically in dorsal hippocampus (Leontiadis et al., 2023). Parvalbumin-expressing inhibitory interneurons in the hippocampus are important for pacing pyramidal cell spiking during sharp wave-ripples (Schlingloff et al., 2014; Stark et al., 2014), and hippocampal network models have suggested that inhibition within CA1 is important for controlling cell participation within replay events (Ramirez-Villegas et al., 2018). Reported weaknesses in pyramidal cell-interneuron coupling in CA1 have only been examined during active behavior in FXS models (Arbab, Battaglia, et al., 2018). However, considering that ripple oscillations are locally generated in CA1 (Csicsvari et al., 1999), the lower peak ripple frequencies that we found during replay events may suggest that firing of CA1 inhibitory interneurons is also slowed during ripples in FXS rats. Due to the limited number of interneurons in our dataset, we were not able to test this hypothesis directly in the present study.

Both CA3 and CA2 contribute to generation of sharp wave-ripples and replay events in CA1 (Csicsvari et al., 1999, 2000; Oliva et al., 2016). Thus, deficits in CA1 place cell firing in FXS rats during replay events may be inherited from these upstream regions or due to local deficits in CA1. Future studies involving simultaneous recordings from CA3, CA2, and CA1 are necessary to shed light on circuit mechanisms underlying slowed firing of place cells during replay events in FXS rats.

In addition to inputs to CA1 from CA3 and CA2, input from the medial entorhinal cortex (MEC) to CA1 can be important for replay events, particularly replay events that span multiple sharp wave-ripples (Yamamoto & Tonegawa, 2017). Previous work has suggested that synaptic inputs from MEC to CA1 pyramidal cells are reduced in FXS models (Asiminas et al., 2022; Ordemann et al., 2021). Although we did not observe a difference in the number of ripples co-occurring with replay events of a given duration between FXS and WT rats (Figure 8E), a reduction in MEC input during replay events may affect temporal compression of replayed sequences for extended replay events in CA1.

Downstream targets of the hippocampus may be affected by impaired temporal compression of awake replay events. During replay events, activity between hippocampus and prefrontal cortex is coordinated (Berners-Lee et al., 2021; Harvey et al., 2023; Jadhav et al., 2016; Peyrache et al., 2009; Shin et al., 2019; Tang et al., 2017). This coordinated activity has been hypothesized to support memory retrieval processes that can be used to guide future decision making (Jadhav et al., 2016; Zielinski et al., 2020). The strength of excitatory drive from the hippocampus to the prefrontal cortex during sharp wave-ripples can affect the response of the prefrontal cortex (Wierzynski et al., 2009), suggesting that abnormally slowed spike timing during replay events in CA1 of FXS rats may alter subsequent prefrontal responses.

The work presented shows novel evidence for specific physiological impairments in the hippocampus in a rat model of Fragile X Syndrome. However, these impairments were characterized using a simple behavioral protocol with no memory component. Rats were familiarized to the environment before recording, and food rewards were randomly given without any motivational salience for specific spatial trajectories. Previous work suggests that the temporal compression of place cell sequences during both active behavior and awake rest can contribute to spatial learning and memory. The slopes and strength of theta sequences have been reported to increase during learning of new environments or trajectories to a new goal location (Feng et al., 2015; Igata et al., 2021; Pfeiffer, 2022), and theta sequences exhibited higher slopes during correct trials than error trials of a spatial memory task (Zheng et al., 2021). Replay duration and temporal compression have been linked to learning and memory in studies of healthy WT rats (Berners-Lee et al., 2022; Fernández-Ruiz et al., 2019). Replay duration increased, and temporal compression of replay events decreased, across the first several paths that rats took during learning of a new environment (Berners-Lee et al., 2022). However, on a longer time scale, the duration of replay events decreased, while the lengths of trajectory representations increased, across sessions in rats learning a spatial memory task (Shin et al., 2019). These results raise the possibility that less temporally compressed theta sequences and slow replay could contribute to impaired spatial learning and memory in FXS rats. Future studies of place cell population activity in FXS rats engaged in learning and memory tasks will be important to shed light on this question.

## Acknowledgements

This research was supported by the Department of Defense CDMRP award W81XWH1810314 (to L.L.C.) and National Institutes of Health awards R56MH125655 (to L.L.C), R01MH131317 (to L.L.C.), and F31MH127933 (to M.M.D.). The authors thank Isabella Lee, Misty Hill, Ayomide Akinsooto, and Sirisha Dhavala for technical assistance and Chenguang Zheng, Ernie Hwaun, and John Trimper for providing MATLAB code for some of the analyses. The authors acknowledge the Texas Advanced Computing Center (TACC) at The University of Texas at Austin for providing data storage resources that have contributed to the research described within this article. URL: http://www.tacc.utexas.edu

## Author Contributions

M.M.D. and L.L.C. designed research; M.M.D. and E.R. performed research; M.M.D. analyzed data; M.M.D. and L.L.C. wrote the paper.

